# Direct NMR evidence for transient binding of the vesicular SNARE protein synaptobrevin-2/VAMP2 to the α-synuclein C-terminus

**DOI:** 10.64898/2026.11.03.764002

**Authors:** Sophia Werner, Tobias Stief, Celina M. Schulz, Katharina Vormann, Robin Backer, Marie P. Schützmann, Lothar Gremer, Wolfgang Hoyer, Nils-Alexander Lakomek

## Abstract

The protein α-synuclein (α-syn) plays a central role in several neurodegenerative diseases, including Parkinson’s disease; however, its physiological function remains enigmatic. We provide direct evidence by nuclear magnetic resonance (NMR) spectroscopy, microscale thermophoresis (MST), and biolayer interferometry (BLI) measurements that the α-synuclein C-terminus binds to the vesicular SNARE (soluble N-ethylmaleimide-sensitive factor attachment protein receptor) protein synaptobrevin-2 (syb-2), also known as VAMP2 (vesicle-associated membrane protein). The observed dissociation constant (Kd) in the 10–50 micromolar range falls within the estimated concentration range of α-synuclein in neurons, while local concentrations may be substantially higher. NMR chemical shift perturbation (CSP) and paramagnetic relaxation enhancement (PRE) experiments mapped the binding site with residue-specific resolution to the α-synuclein C-terminus (V118–E137) and to the synaptobrevin-2 SNARE motif/linker domain interface (region A82–K87), and also affected the synaptobrevin-2 N-terminus (M1–S2). NMR CSPs and PREs indicate a transient interaction in the fast exchange (µs-ms) regime. These transient weak binding interactions are characteristic of intrinsically disordered proteins (IDPs) and allow both proteins to engage in multiple binding interactions, thereby enabling α-synuclein to play a regulatory role upstream of or during neuronal exocytosis. While the exact physiological role of α-synuclein remains debated, the involvement of its pathological form in several neurodegenerative diseases (dubbed synucleinopathies) is well established. Our data suggest a direct link between proteins involved in neurodegeneration and neuronal exocytosis.

## Introduction

The protein α-synuclein (α-syn) is located predominantly in the presynaptic terminals of neurons^1^. Since a high abundance of α-syn was detected in Lewy body deposits extracted from post-mortem brain samples of patients with Parkinson’s disease or Lewy body-related Alzheimer’s disease^2^, the pathological form of α-syn has stirred substantial biochemical research related to these neurodegenerative processes. The plethora of neurodegenerative diseases associated with α-syn misfolding has been dubbed *synucleinopathies*^3–5^.

The protein α-syn belongs to the class of intrinsically disordered proteins (IDP)^6–9^. However, under certain conditions, α-syn can adopt structured conformations. so that it can exist in a soluble, disordered state and a partially structured, membrane-bound state. Via its amphipathic N-terminal domain (NTD, residues M1–K60; for the α-syn domain scheme, see **Figure 1a** and **Figure S1a** for the amino acid sequence), the protein associates with cellular membranes, where it adopts an α-helical structure^10–14^. Alternatively, in the pathological case, α-syn can form toxic oligomers and ultimately amyloid fibrils, which have both been linked to *synucleinopathies*^3–5,15,16^. Here, the NAC (non-amyloid β-containing) domain, also known as the hydrophobic region (HR)^17^, is the central region, participating in fibril formation by adopting β-sheet secondary structure, whereas the highly acidic C-terminal domain (CTD) remains intrinsically disordered (**Figure 1a**). Although the structure of these amyloid fibrils has been extensively characterized, the exact pathological mechanisms underlying neurodegeneration remain not well understood.

**Figure 1:**
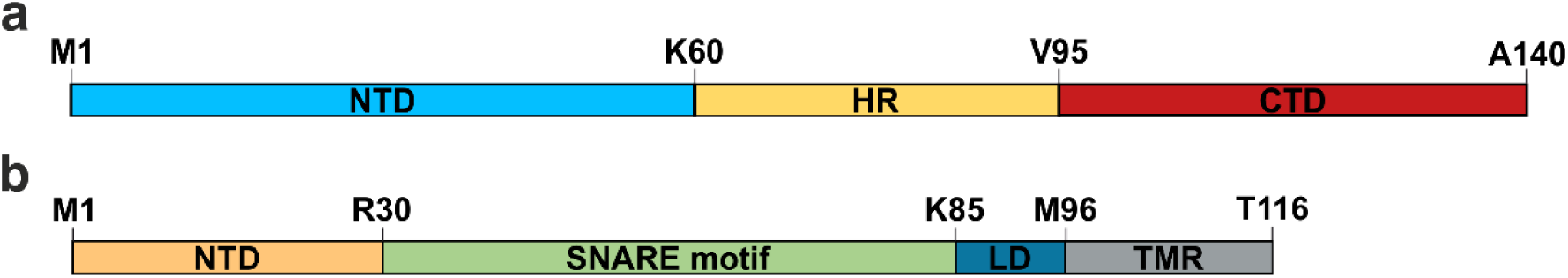
Schematic representation of α-synuclein (a) and synaptobrevin-2 (b). (**a)** Domain scheme of α-syn with the N-terminal domain (NTD, M1–K60) highlighted in light blue, the hydrophobic region (HR, E61 to V95) in yellow, and the C-terminal domain (CTD, K96–A140) in red. (**b)** Domain scheme of syb-2, with the N-terminal domain (NTD, M1–N29) highlighted in orange, the SNARE motif (R30– K85) in green, the linker domain (LD, R86–M95) in blue, and the C-terminal transmembrane region (TMR, M96–T116) in grey. In the current study, we focused on the cytoplasmic part of syb-2, syb-2(1– 96), without the TMR.

Even less is known about the physiological function of α-syn. One of the first proposed physiological roles of α-syn was a non-classical chaperone activity during neurotransmitter release^18^. Neurotransmitter release occurs rapidly and at high frequency at presynaptic nerve terminals, usually at sites distant from the neuronal cell body. It requires frequent cycles of SNARE (soluble N-ethylmaleimide-sensitive factor attachment protein receptor) complex assembly. In their monomeric form, SNARE proteins are intrinsically disordered^18–22^. Burré and Südhof found that α-syn interacts with the vesicular SNARE protein synaptobrevin-2 (syb-2)^18^, also known as vesicle-associated membrane protein 2 (VAMP2) (**Figure 1b** and **Figure S1b**). Using a combination of immunoprecipitation and biochemical truncation assays, an interaction between the α-syn C-terminus and the syb-2 N-terminus was suggested^18^. The interaction between α-syn and syb-2 in the context of SNARE complex assembly was later refined to specifically involve the membrane-bound form of α-syn, with membrane-bound α-syn multimers serving as chaperones for SNARE complex assembly^23^. Lou et al. proposed that α-syn acts as a cross-bridge between syb-2 and acidic phospholipids, thereby facilitating SNARE-dependent vesicle docking^24,25^. C-terminal truncation of α-syn abrogated this effect. However, it remains controversial whether α-syn facilitates^24,25^ or reduces SNARE-dependent neurotransmitter release^26–28^. Alternative physiological roles of α-syn, independent of SNARE complex formation, have been proposed, particularly in the regulation of synaptic vesicle clustering and storage near the active zone, upstream of vesicle docking^29^. This concept was recently extended by linking α-syn to the maintenance of biomolecular condensates associated with the synaptic vesicle pool^30,31^. The exact molecular function of α-syn, however, remains elusive and highly debated^32–35^.

Here, we investigated the interaction between α-syn and syb-2 using NMR spectroscopy, microscale thermophoresis (MST), and biolayer interferometry (BLI). We established a Kd in the low µM range. While the syb-2 linker domain is the primary interaction site, the syb-2 N-terminus also appeared to be affected by the binding interaction. We assigned the binding epitope to V118–E137 on the α-syn side and to the positively charged interface of the C-terminal SNARE motif and linker domain region A82–K87 on the syb-2 side, with residue-specific resolution.

## Results

To investigate the interaction between syb-2(1–96) and α-syn, we performed MST and BLI measurements, NMR titration, and Paramagnetic Relaxation Enhancement (PRE) experiments. MST measurements revealed an interaction between syb-2(1–96) and α-syn, with a dissociation constant of Kd of 14 ± 4 µM (**Figures 2a** and **2b**). Using BLI measurements, a Kd of 45.3 ± 19.3 µM was detected (**Figures 2c** and **2d**). Therefore, a weak binding interaction between α-syn and syb-2(1–96), with affinities in the low-micromolar range, was consistently observed by both independent methods.

**Figure 2:**
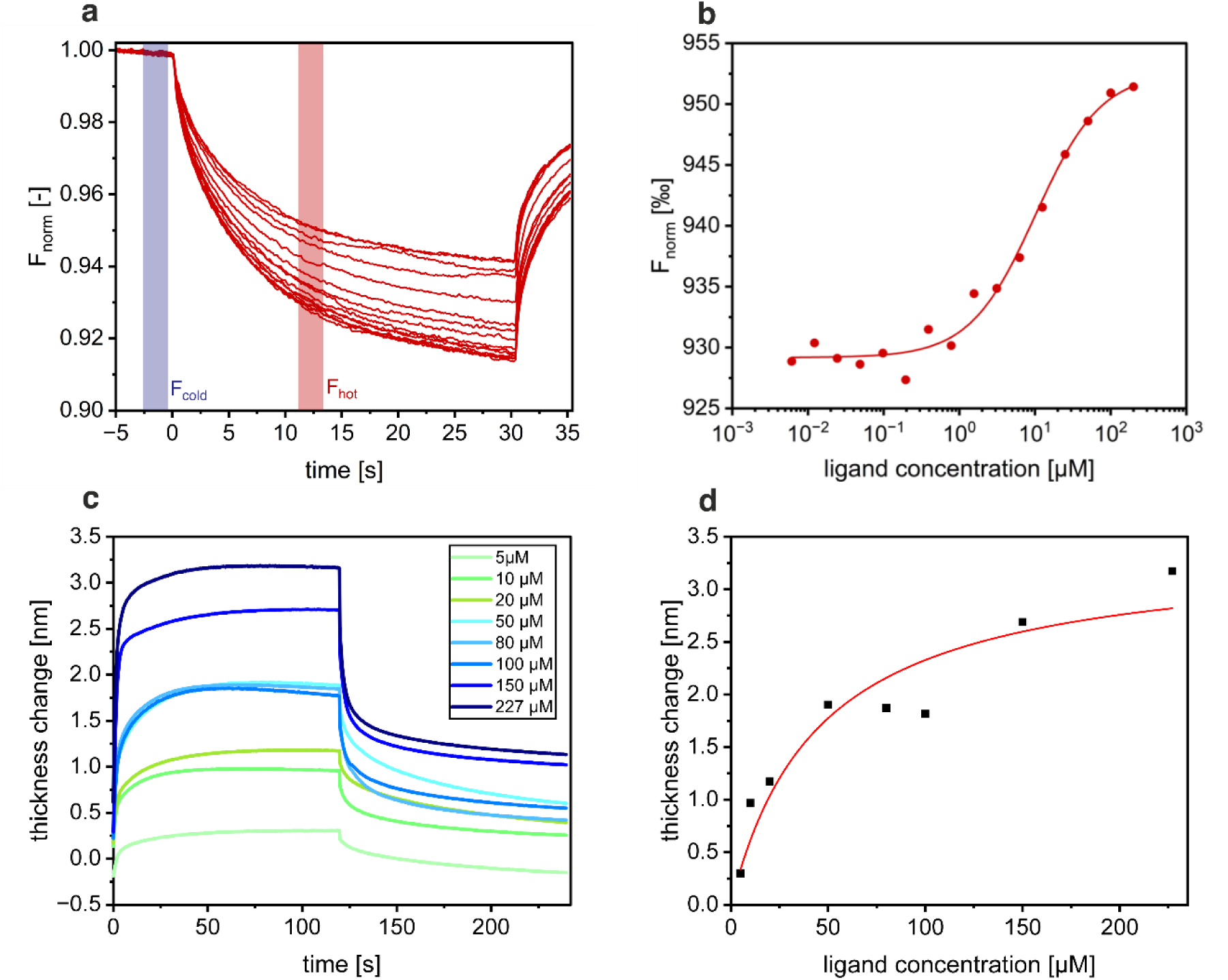
Binding interaction between α-synuclein and synaptobrevin-2, determined by (a, b) MST and (c, d) BLI measurements. **(a)** The MST traces of the relative fluorescence of α-syn-Alexa488 were plotted against time. F_cold_ describes the homogeneous fluorescence distribution when the infrared (IR) laser is off. F_hot_ describes the alternate movement across the temperature gradient when the IR laser is on, resulting in a binding-dependent drop in fluorescence. **(b)** Normalized fluorescence (F_norm_) in ‰ against the ligand concentration (syb-2(1–96)), plotted on a logarithmic scale. A 1:1 binding model was fitted, yielding an equilibrium constant of K_d_ of 14 ± 4 µM. **(c)** Biotinylated and sensor-bound α-syn is exposed to an increasing excess concentration of syb-2(1–96). The change in biolayer thickness resulting from syb-2(1–96) binding is plotted over time. The color gradient represents the concentrations of syb-2 (1–96) added. In **(d),** the mean plateau value of the thickness change between 60 s and 120 s was plotted against the ligand concentration. The data were then fitted using **Equation 1**.

NMR titration experiments were conducted to gain residue-specific insights into the interaction of α-syn and syb-2(1–96) (**Figure 3**). Starting with ^15^N-labeled α-syn (visible in 2D NMR ^1^H– ^15^N HSQC spectra), we performed an NMR titration series by adding an excess of unlabeled syb-2(1–96) (invisible in 2D ^1^H–^15^N NMR spectra) to ^15^N-labeled α-syn (NMR visible), in a stoichiometric molar ratio ranging from 1:1 to 1:5 of α-syn:syb-2(1–96). While most NMR resonances in the 2D ^15^N HSQC spectrum of ^15^N-labeled α-syn remained unchanged upon the addition of unlabeled syb-2(1–96), distinct chemical shift perturbations (CSPs) were observed for the region V118–E137 compared to the reference spectrum without syb-2(1–96). The CSPs increased linearly with higher molar excess of syb-2(1–96), up to the tested stoichiometric ratio of α-syn to syb-2(1–96) of 1:5, both at 20 °C (**Figures 3a, 4a,** and **S3)** and 37 °C **(Figures 4c, S3,** and **S4a**). An exception was residue H50, which also exhibited chemical shift changes but without a clear trend. The largest CSP upon syb-2(1–96) binding was detected for Y125. To confirm that the α-syn C-terminus is indeed the binding site, we recorded the MST measurements with a C-terminally truncated α-syn mutant (α-syn^Δ^ ^Q109–A140^). Indeed, binding was substantially reduced for the truncation mutant, rendering reliable Kd estimation impossible (**Figure S2**).

**Figure 3:**
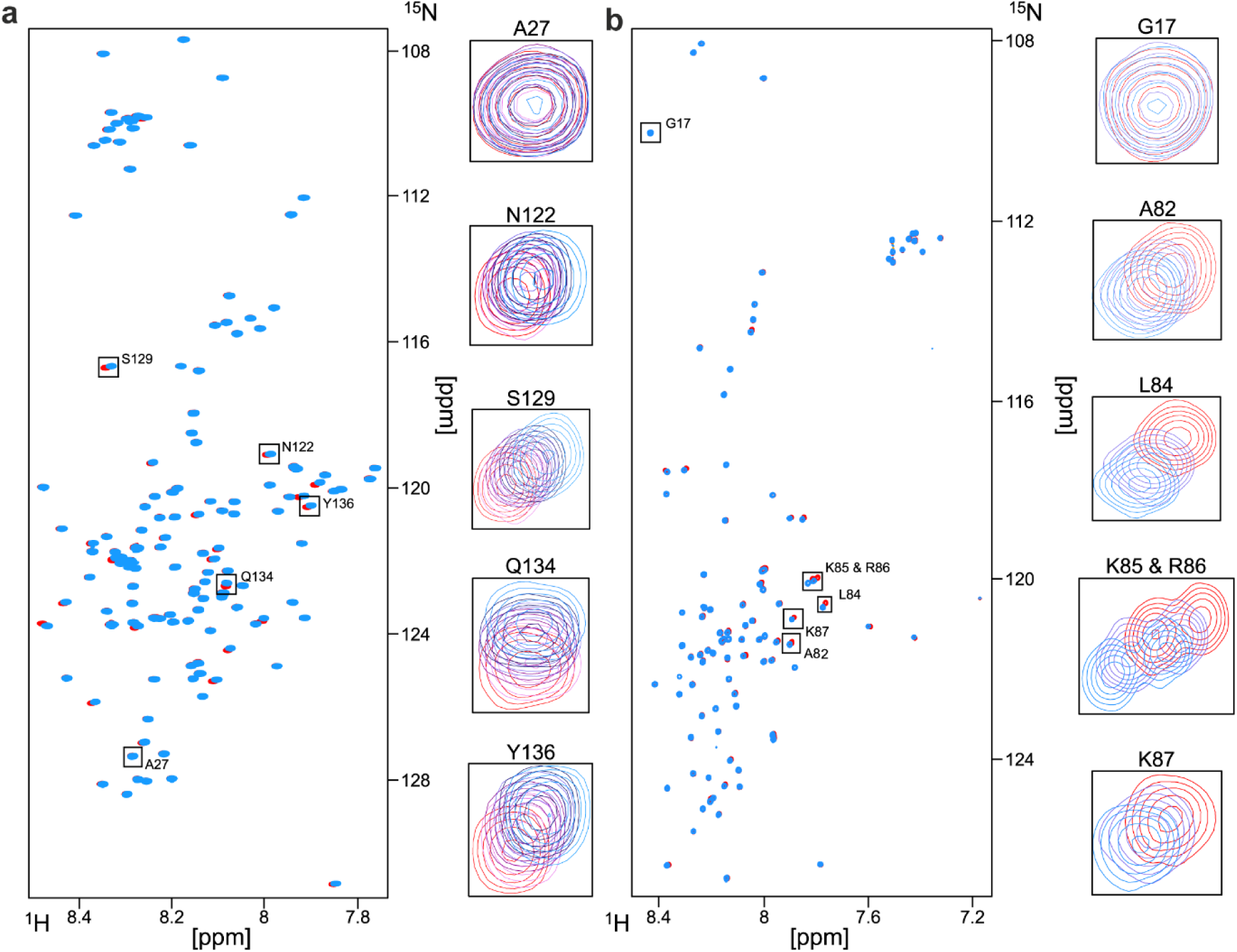
NMR titration of α-synuclein with synaptobrevin-2(1–96) and vice versa. Overlay of 2D HSQC spectra of NMR visible 40 µM ^15^N α-syn in the presence of syb-2(1–96) (1:5, blue) and in the absence of syb-2(1–96) (red) **(a**) and further, of 40 µM ^15^N syb-2(1–96) in the presence of α-syn (1:5, blue) and in the absence of α-syn (red) (**b**), measured at 1.2 GHz and 20 °C. Representative residues with significant chemical shifts are shown separately with an overlay of all titration steps (reference, red; and molar ratios of 1:1, pink; 1:2, light violet; 1:3, dark violet; 1:4, dark blue; 1:5, light blue), as well as one example of no significant chemical shift, for reference (α-syn A27 and syb-2(1–96) G17, respectively).

**Figure 4:**
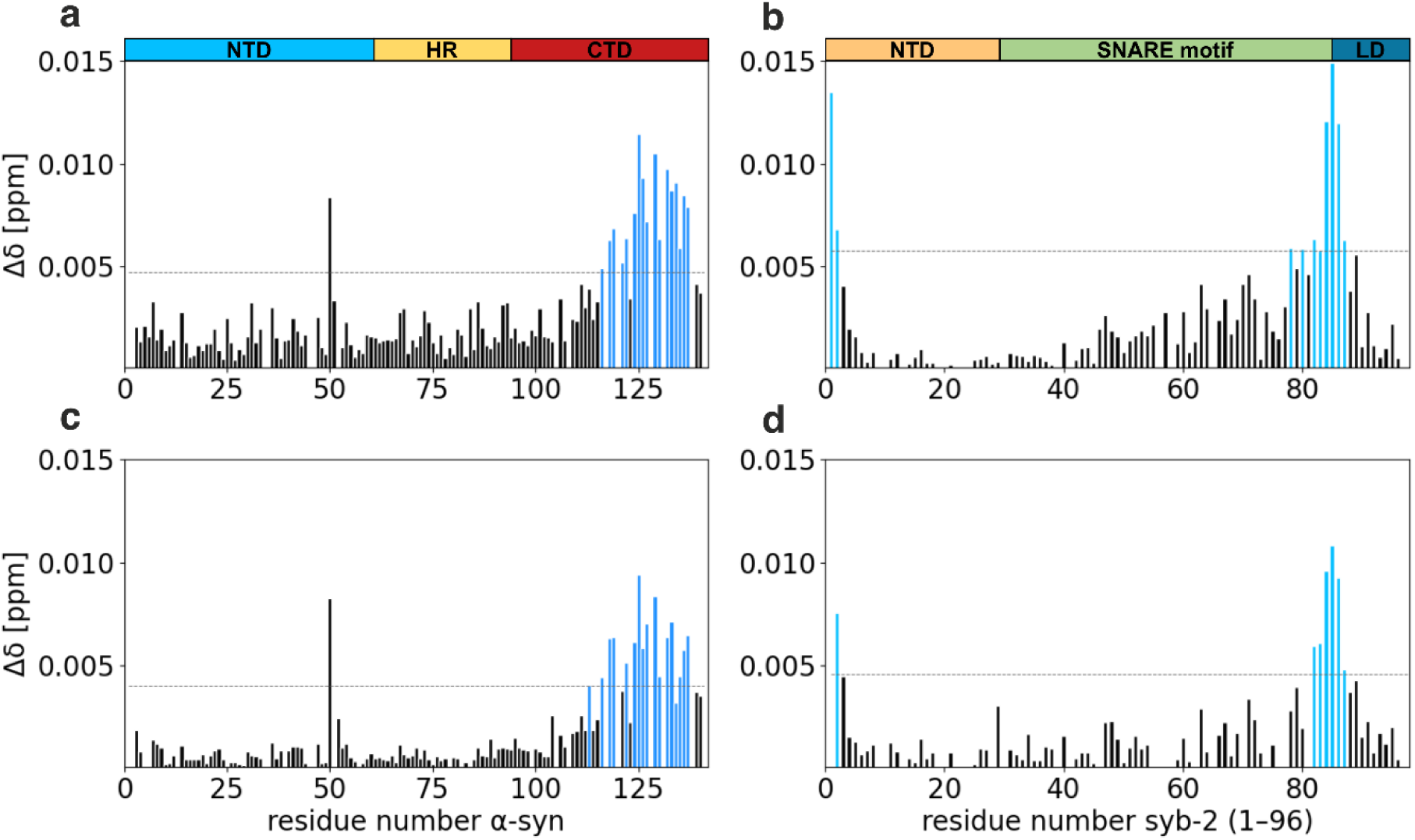
NMR chemical shift perturbation analysis of the α-synuclein/synaptobrevin-2(1–96) binding. CSP (Δδ) of α-syn upon titration of syb-2(1–96) (**a** and **c**) and vice versa (**b** and **d**). CSPs exceeding two times the standard deviation (grey line) of the CSP distribution upon binding (2*STEDV) are highlighted in dark blue (α-syn) and in light blue (syb-2(1–96)). **a** and **c** show the chemical shift changes (Δδ) for the ^15^N α-syn to syb-2(1–96) ratio of 1:5, relative to the ^15^N α-syn reference without syb-2(1–96) at (**a)** 20 °C and (**c)** 37 °C. **(b)** and **(d)** show the CSPs (Δδ) for the ratio 1:5 of ^15^N synaptobrevin-2(1–96) to α-syn at (**b**) 20 °C and (**d)** 37 °C. The domain schemes of α-syn and syb-2(1– 96) (cf. Figure 1) are plotted on top.

To obtain further insights into the binding epitope on the syb-2(1–96) side and monitor the binding interaction from this perspective, we performed an analogous titration series with ^15^N-labeled syb-2(1–96) (NMR visible) and unlabeled α-syn (NMR invisible) (**Figure 3b**). Similar to α-syn, syb-2(1–96) also exhibits a narrow chemical shift dispersion in the ^1^H dimension, characteristic of an IDP. The chemical shifts of the majority of resonances remained unchanged upon the addition of α-syn. Still, characteristic chemical shift changes were observed for the C-terminal SNARE motif (R30–K85) and the linker domain (R86–M96) of syb-2(1–96), most prominent for the region A82–K87 centered around K85 (**Figure 4b)**, at the interface between the SNARE motif and linker domain (**Figure 1b**). In addition, we observed chemical shift changes for residues M1 and S2 (**Figure 3b** and **4b**). These characteristic chemical shift changes were detected both at 20 °C (**Figure 3b** and **4b**) and 37 °C (**Figure 4d, S3,** and **S4b).**

To confirm the binding of the α-syn C-terminus to the syb-2(1–96) linker domain, we attached MTSL ((1-Oxyl-2,2,5,5-tetramethyl-δ-3-pyrroline-3-methyl) methanethiosulfonate), a nitroxide radical spin label (paramagnetic), to an artificial Cysteine residue at the α-syn C-terminus (α-syn^A140C^) and conducted NMR paramagnetic relaxation enhancement (PRE) experiments. The paramagnetic enhancement of the ^1^H R2 relaxation rate constants was measured for ^15^N syb-2(1–96). Regions of syb-2(1–96) close to the α-syn C-terminus (and the paramagnetic spin-label) will show increased ^1^H R2 rate constants. We found the most substantial ^1^H R2 increase for the syb-2(1–96) linker domain region, confirming the interaction between the α-syn C-terminus and the syb-2(1–96) linker domain (**Figure 5**). The paramagnetic enhancement of the ^1^H R2 rate constants scales with r^-6^, where r is the distance to the spin label. In addition to the linker domain region, a weak increase of the ^1^H R2 rate constants was observed for the entire SNARE motif (residues R30–S80), indicating that also the syb-2 SNARE motif is in proximity to the α-syn C-terminus, albeit more distant than the syb-2(1–96) linker domain. Interestingly, the syb-2(1–96) N-terminus does not experience any PRE effect.

**Figure 5:**
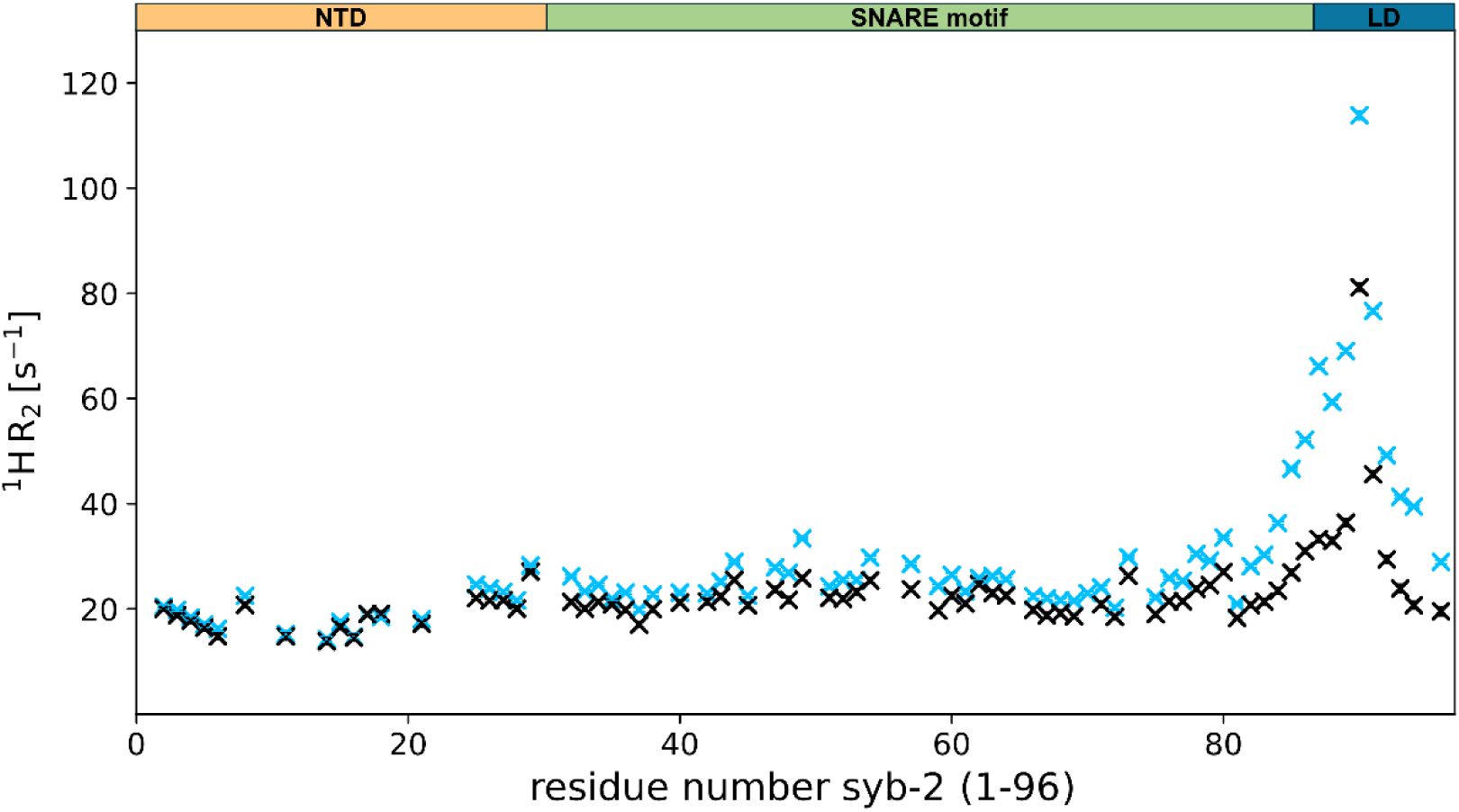
Paramagnetic relaxation enhancement measurement. The ^1^H R_2_ rate constants of ^15^N syb-2(1–96) were plotted against the residue number in the presence of α-syn with the C-terminal attached nitroxide paramagnetic spin label (light blue) and compared to the rate constants of the diamagnetic reference sample with quenched spin label (black).

## Discussion

For the presynaptic protein α-syn, the pathological link to Parkinson’s disease and Lewy-body-associated dementia has been established^2–5^. Nevertheless, the physiological role of α-syn remains largely unclear^32–35^. While we refrain from pinpointing a primary function of α-syn in its native form, which may fulfill multiple physiological roles, we hereby establish a transient binding interaction between α-syn and syb-2 and identify the α-syn C-terminus and the syb-2 SNARE motif/linker domain interface as the primary binding epitope. Consequently, potential physiological roles of α-syn in synaptic vesicle storage or trafficking appear reasonable^29–31^, as well as during SNARE-dependent vesicle fusion^18,23–28^.

BLI, MST, and NMR measurements consistently indicate weak binding between α-syn and syb-2(1–96), with a Kd in the 10–50 μM range. The Kd is within the physiological concentration range of α-syn found in neurons^36^. NMR CSP data, recorded during NMR titration series, map the binding interface to the α-syn C-terminus (V118–E137) and the syb-2 SNARE motif C-terminus/linker domain interface (A82–K87). Furthermore, chemical shift changes are observed in the two N-terminal syb-2 resonances. **Figure 6** displays the electrostatic surface of α-syn and syb-2(1–96). Both proteins show a high percentage of charged acidic and basic residues, as expected for IDPs. While the α-syn C-terminus is highly negatively charged, the syb-2 linker domain is of opposite positive charge. Therefore, it is reasonable that both regions will attract each other, suggesting that electrostatic interactions are a major driving force behind the observed weak interaction between α-syn and syb-2(1–96).

**Figure 6:**
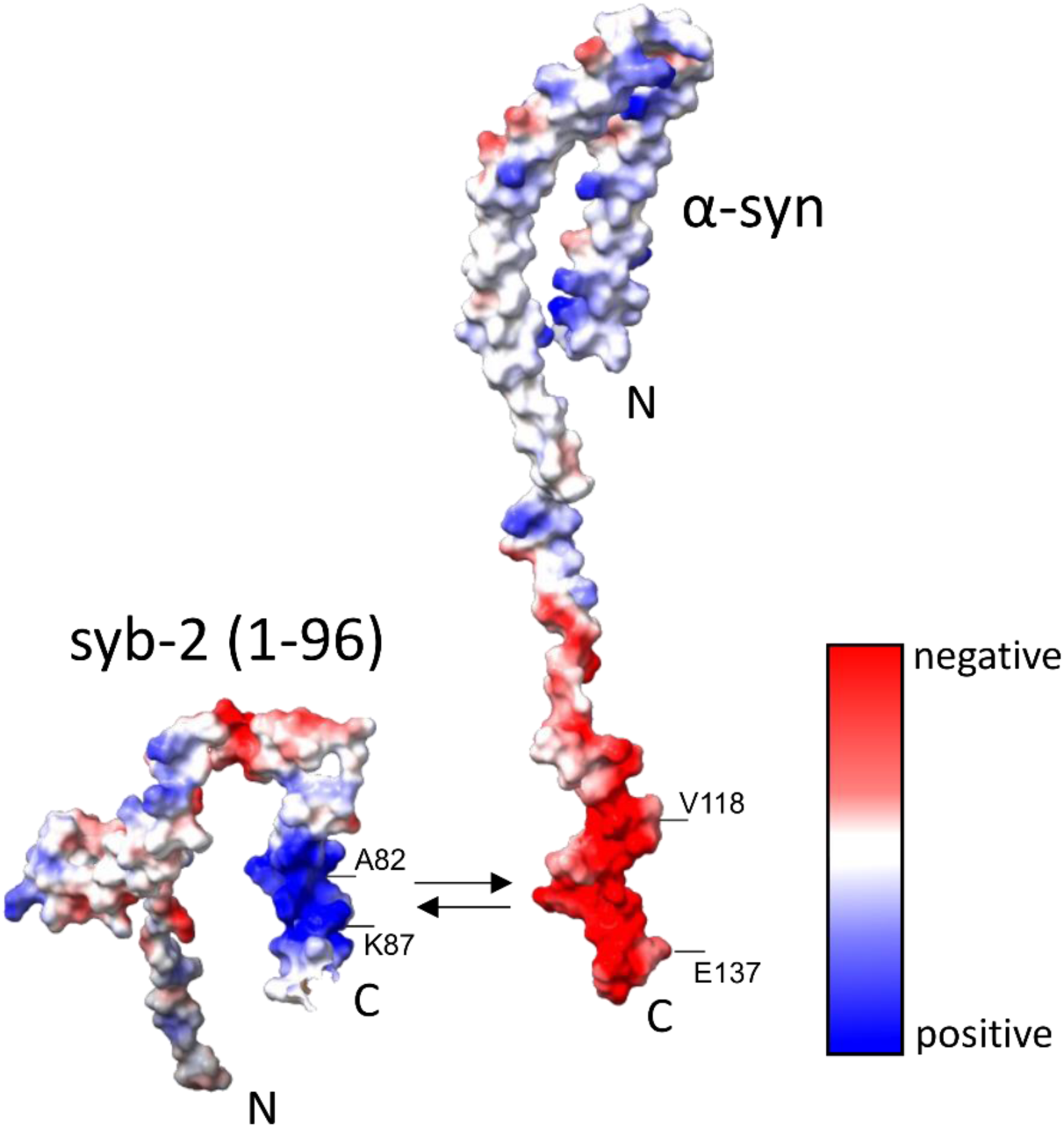
Binding epitope of α-synuclein/synaptobrevin-2 encounter complex. Electrostatic surface potentials of the two proteins, α-syn and syb-2(1–96) are illustrated, with color coding indicating the distribution of electrostatic charges. The color key on the right implies negative (red) and positive (blue) surface charges. Both C-termini (linker domain from syb-2(1–96)) are aligned according to their suggested interaction site [PDB:1XQ8 (α-syn), 2KOG (syb-2(1–96)), created using ChimeraX-1.6.1]^37^. The binding epitopes of α-syn and syb-2(1–96), respectively, identified in this work are highlighted on the right side of each protein. For α-syn, residues D119, D121, E123, E126, E130, E131, D135, and E137 contribute to the negative surface charge of the binding region. For syb-2(1–96), the positive surface charge is primarily contributed by residues K83, K85, R86, and K87.

The C-terminus of α-syn is known to be intrinsically disordered and does neither participate in α-syn membrane binding^6,10–13^ nor in amyloid fibril formation^3–5,15,16^. For the syb-2 linker domain and C-terminal SNARE motif, we recently found that it transiently binds to lipid membranes, leading to the disappearance of signal intensities for the linker domain residues as well as the C-terminal SNARE motif with a Kd in the hundreds of micromolar range^19^. To probe a potential interaction between the syb-2 linker domain and α-syn, we have refrained from C-terminally membrane-bound full-length syb-2 in this study and focused on the soluble part of syb-2, namely syb-2(1–96). Probing interactions in the context of lipid bilayers and full-length syb-2(1–116) will be the scope of future research.

NMR PRE measurements employ a covalently attached paramagnetic spin-label that increases the ^1^H relaxation rate constants of nearby residues. PRE data of C-terminally labeled α-syn indicate transient proximity between the α-syn C-terminal domain and the syb-2 linker domain, confirming the binding region as revealed by the NMR CSP data. In addition to the syb-2 linker domain, two N-terminal syb-2 residues (M1 and S2) also show differences in the CSPs upon addition of α-syn. The syb-2 N-terminus was previously suggested as the main site of interaction with α-syn^18^. In contrast to the CSPs, the NMR PRE data do not show any effect on the syb-2 N-terminus.

Recently, two independent studies monitored syb-2-driven phase separation of α-syn^30,31^. Wang et al. proposed that syb-2 acts as a chaperone for α-syn in synaptic vesicle co-condensates^31^. Both studies reported NMR CSPs of the α-syn C-terminus in the presence of syb-2^31^ or a peptide mimicking the syb-2 isolated linker domain^30^. In addition, Wang et al. found distinct CSPs for the region S80-K87^31^, in good agreement with the binding epitope identified by our NMR study. For α-syn, the binding epitope reported by Wang et al. was less specific but also encompassed the binding region identified here, V118–E137. This further narrows the initially suggested interaction region from A95–A140^18^ towards V118–E137 and challenges a previously reported interaction site V96–E110^26^. Moreover, co-condensates composed of α-syn and syb-2 with protein-free synaptic vesicle mimics were visualized by Wang et al.^31^. Disruption of the α-syn/syb-2 interaction by C-terminal truncation of α-syn abolished the observed co-condensation^31^.

Beyond the study by Wang et al., our extensive biomolecular interaction data, measured by MST and BLI, consistently yield Kd values in the low micromolar range. A Kd in the micromolar range was previously reported by Fakhree et al.^38^ as well. As a negative control, MST measurements with a C-terminally truncated variant of α-synuclein (α-synuclein^ΔQ109–A1^^40^), lacking the last 30 amino acids, showed a markedly diminished binding affinity.

Our PRE data support the conclusion that the α-syn/syb-2 interaction extends beyond the immediate binding interface. Furthermore, NMR CSP data revealed that the syb-2 N-terminus is influenced by α-syn binding.

Both α-syn and monomeric syb-2 belong to the essential but under-investigated class of IDPs. With about 25% of the human proteome being IDPs and 40% of the human proteome being proteins with large intrinsically disordered regions (IDRs) (including IDPs, thus about 60% of proteins being folded in the classical sense)^39^. In the context of neuronal exocytosis, we rationalize the transient nature of the α-syn/syb-2 interaction by a potential regulatory role of α-syn upstream or during neuronal exocytosis. A cascade of syb-2 interactions with increasing affinity of the binding partner would tightly regulate and orchestrate SNARE complex assembly and subsequent synaptic vesicle fusion. In line with vesicle storage or vesicle docking hypotheses, a low-affinity interaction between α-syn and syb-2 might allow α-syn to modulate syb-2 function without interfering with the SNARE complex.

In this study, we have demonstrated, at the molecular level with residue-specific NMR resolution, a transient binding interaction between α-syn and the SNARE protein syb-2, which participates in SNARE complex formation and drives neuronal exocytosis. Consequently, this would be consistent with the loss-of-function hypothesis for aggregated α-syn, which contributes to the pathogenesis of neurodegenerative diseases. Misfolded and aggregated α-syn would thus be unable to fulfill its yet-to-be-fully-characterized physiological function. Interestingly, phosphorylation of Y125 and S129, which are both part of the syb-2 binding site, as well as C-terminal truncation of α-syn, seem to be connected to the pathological function of α-syn^40–44^.

To summarize, using MST and BLI measurements, we consistently identified a transient binding interaction between α-syn and the vesicular SNARE protein syb-2 in the 10–50 µM range. Both proteins are intrinsically disordered in their monomeric form and remain so upon interaction, as indicated by the NMR CSP data. Two distinct binding sites were identified: the negatively charged region V118–E137 at the intrinsically disordered C-terminus of α-syn binds the positively charged linker domain region A82–K87 on the syb-2 side (**Figure 6**). The opposite charges in the two regions indicate that electrostatic interactions are the primary driving force behind the weak, transient binding.

## Methods

### Protein sample preparation

The heterologous expression in *E.coli* and purification of synaptobrevin-2(1–96) (origin: *Rattus norvegicus*) were performed as described previously (96 amino acids plus three N-terminal residues (GSH), 10.78 kDa, pI = 8.28, extinction coefficient ε280nm = 12490 M^-1^ cm^-1^, see Figure **S1** for sequence information)^19,45,46^. α-synuclein wildtype (origin: *Homo sapiens,* 140 amino acids, 14.46 kDa, pI = 4.67, extinction coefficient ε275 nm = 5600 M^-1^ cm^-1^), α-syn^A140C^ and α-syn^Δ^ ^Q109–A1^^40^ (ε275 nm = 1400 M^-1^cm^-1^) were expressed and purified as described previously^47,16^. ^15^N-labeled samples for NMR measurements were grown in minimal medium (M9) supplemented with ^15^N NH4Cl (99 %, Cambridge Isotope Laboratories, USA).

For PRE measurements, α-syn^A140C^ was paramagnetically labeled with (1-Oxyl-2,2,5,5-tetramethyl-δ-3-pyrroline-3-methyl) methanethiosulfonate (MTSL or MTSSL, Toronto Research Chemicals, O875000). For labeling, MTSL was dissolved in 90 µL DMF (dimethylformamide) and 810 µL 200 mM HEPES (2-(4-(2-Hydroxyethyl)-1-piperazinyl)-ethanesulfonic acid), pH 7.4, and added to lyophilized α-syn^A140C^. Labeling was performed with a five-fold molar excess of MTSL for 1 h at room temperature (∼20 °C). For purification, reversed-phase HPLC was performed on an Agilent Infinity 1260 HPLC system using a semi-preparative ZORBAX 300SB-C8 column (9.6 x 250 mm, Agilent, 880995-206). The purification was carried out using an acetonitrile (ACN) gradient, ranging from 8% (v/v) ACN to 80% ACN, in aqueous 0.1% (v/v) trifluoroacetic acid (TFA), at 40 °C, over 40 min, and with a flow rate of 4 mL/min.

Protein concentrations were determined by measuring the UV absorbance at 275 nm (α-syn) or 280 nm (syb-2(1–96)), respectively. The protein samples were flash-frozen in 20 mM sodium phosphate, pH 6.0, 50 mM NaCl, and stored at-80 °C until use, except for MTSL-labeled α-synA140C, which was lyophilized overnight in a SpeedVac (Christ LT-105, Christ RVC 2-18) after labeling and stored at-20 °C.

### Biolayer interferometry (BLI)

BLI measurements were performed using N-terminal biotinylated α-syn bound to streptavidin-coated biosensors, “High precision Streptavidin (SAX), Dip and Read Biosensors” (fortéBIO) and conducted on the BLItz instrument by fortéBIO (Sartorius).

First, the biosensor was hydrated in the sample buffer (20 mM sodium phosphate, pH 6.0, 50 mM NaCl). A baseline was recorded for 30 seconds. The streptavidin sensor was loaded with 1.15 μM N-terminal biotinylated α-syn for 250 seconds. A second baseline was recorded for 30 seconds, followed by a saturation step in which 1 mM biotin was applied for 60 seconds. This step ensures that syb-2(1–96) does not interact with the biosensor. A baseline was recorded for 30 seconds. The association of the analyte syb-2(1–96) was probed by applying increasing concentrations of the analyte (5 µM, 10 µM, 20 µM, 50 µM, 80 µM, 100 µM, 150 µM and 227 µM). After recording the association for 120 seconds, the sensor was dipped in buffer, and dissociation of syb-2(1–96) was recorded for 120 seconds. The program *Advanced Kinetics* was used to operate the BLI instrument. For analysis, an equation analogous to the one describing enzyme saturation was used to fit the plateau value of the binding curve and the dissociation constant (**Equation 1**). A 1:1 binding model was assumed. For the mean plateau value of the thickness change Y, the data range was 60–120 s. The plateau value Ymax and the dissociation constant Kd were fit parameters.

**Equation 1**: Plateau fit of the binding curve. Y describes the measured thickness change [nm], [L] is the ligand concentration [µM], and K_d_ describes the dissociation constant.

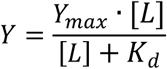

### Microscale thermophoresis (MST)

To determine the dissociation constant (Kd), MST measurements were performed using α-syn wildtype labeled with Alexa Fluor488 NHS-Ester (Thermo Fisher Scientific).

For labelling, Alexa Fluor488 dissolved in DMSO was added to the purified protein at a molar ratio of 1:3 and incubated for 1 h under light-protected conditions on an overhead shaker. To reduce the lysine reactivity and favour the reaction with the N-terminus, we performed the labelling at pH 6.0 (20 mM sodium phosphate, pH 6.0, 50 mM NaCl). Furthermore, only samples with a degree of labelling (DOL) between 50% and 100% were used.

Free Alexa Fluor 488 was subsequently removed using a PD-10 Mini desalting column (PD MiniTrapTM G-25 Sephadex, 2.1 mL, Cytiva). The labelled protein was stored at-20 °C under light-protected conditions. Protein concentration and labelling efficiency were determined by UV–Vis spectroscopy. Absorption spectra were recorded between 250 and 700 nm. Protein absorbance was measured at 275 nm and fluorophore absorbance at 494 nm, and protein concentrations and labelling efficiency (DOL) were calculated using the equations provided below (**Equation 2** and **3**). The molar extinction coefficients of α-syn (ε275 nm) and Alexa Fluor488 (ε494 nm) are 5600 M^-1^ cm^-1^ and 73,000 M^-1^ cm^-1^, respectively. According to the manufacturer, the correction factor (CF) for Alexa Fluor488 NHS-Ester is 0.11. The concentration was calculated with the following equation based on Lambert-Beer’s law.

**Equation 2**: Equation to calculate the protein concentration based on the Lambert Beer Law where c_α-syn_ is the protein concentration [M], A_275 nm_ and A_494 nm_ denote the absorbance at 275 nm and 494 nm, respectively, CF is the correction factor (0.11), ε_α-syn_ is the molar extinction coefficient [M^-1^ cm^-1^], and d is the cuvette path length [cm].

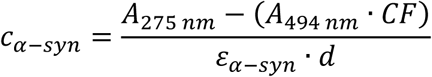

**Equation 3**: The degree of labelling (DOL) was calculated using the equation below, where ε_α-syn_ and ε_Alexa488_ are the molar extinction coefficients (M^-1^ cm^-1^) of α-syn and Alexa Fluor 488, respectively. A_275 nm_ and A_494 nm_ denote the absorbance at 275 nm and 494 nm, respectively. CF is the correction factor (0.11).

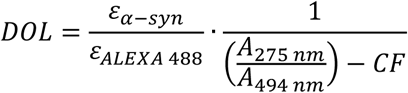

As a ligand syb-2(1–96) was used. The measurements were performed at room temperature (∼24 °C) in 20 mM sodium phosphate, pH 6.0, 50 mM NaCl on a Monolith NT.115 MST instrument (NanoTemper Technologies). The LED intensity and the infrared laser power were set to 50% and 60%, respectively. Starting the measurement, 100 nM α-syn–Alexa488 and 250 nM α-syn^ΔQ109–A140^ Alexa488 were titrated with different ligand (syb-2(1–96)) concentrations up to 200 µM. The samples were transferred to Monolith Premium capillaries. All measurements were performed as triplets. For the C-terminal deletion mutant α-syn^ΔQ109-A140^ experiments were performed analogously. All measurements were analyzed with the MO.Affinity Analysis v2.1.3454 software (NanoTemper Technologies). A 1:1 binding model was fitted according to Seidel et al.^48^.

### Solution NMR spectroscopy

For the titration measurements, 40 µM ^15^N-labeled α-syn or syb-2(1–96) was used. Non-labeled protein was titrated to a concentration of up to 200 µM. The unlabeled protein ligand was added at a molar stoichiometry ratio of 1:1, 1:2, 1:3, 1:4, and 1:5 to ^15^N α-syn. For the measurements with ^15^N syb-2(1–96), the stoichiometry ratios of 1:1, 1:3, and 1:5 were applied. The reference measurement was performed with 40 µM of labeled protein in the absence of non-labeled protein. All measurements were carried out at 20 °C and 37 °C in 20 mM sodium phosphate, pH 6.0, 50 mM NaCl buffer with 5% (v/v) deuterium oxide (D2O, ^1^H2O), 1 mM 3-(trimethylsilyl)-1-propanesulfonic acid sodium salt (DSS), and 0.05% (v/v) NaN3. The titration was measured at a 28 T NMR spectrometer operating at 1.2 GHz ^1^H Larmor frequency (Bruker 1200 MHz AVANCE NEO spectrometer, Bruker Inc., Bellarica), equipped with a 3 mm TCI ^1^H, ^15^N, ^13^C triple resonance cryoprobe. The measurements were processed and analyzed using the Bruker *TopSpin 3.6.* and *CCPN 3 Analysis software*^49^. The NMR resonances of ^15^N α-syn (20 °C) were assigned based on BMRB: 6968 (α-syn). To obtain ^1^H–^15^N assignments at 37 °C, a temperature series of ^1^H–^15^N HSQC spectra in steps of 2 °C was recorded, and the temperature-dependent shifts of resonances were carefully monitored (Supplementary Information Table S1). For syb-2(1–96), NMR assignments (based on the assignments uploaded to the BMRB, accession code 53437) were used and transferred to 20 °C and 37 °C using a temperature series of ^1^H–^15^N HSQC spectra in steps of 2 °C (Supplementary Information **Table S2**). The chemical shift perturbation (CSP) was calculated using the equation shown below (**Equation 4**), whereby all shifts larger than twice the standard deviation were considered significant.

**Equation 4**: Equation for calculating the perturbation of the chemical shift (δ) upon addition of the ligand during the titration. Here, Δδ_av_ is the averaged chemical shift difference, ΔδH is the chemical shift difference in the proton dimension and ΔδN is the chemical shift difference in the nitrogen dimension. The chemical shift differences in the dimensions (ΔδH and ΔδN) were determined as follows^50^.

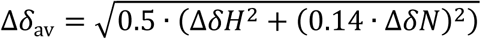

### Statistics

The noise levels of the various NMR spectra were taken as the errors in the NMR spectral intensities. The standard deviation of the NMR chemical shift perturbation (CSP) analysis was calculated assuming a Gaussian normal distribution. The error in the NMR PRE data was calculated using Gaussian error propagation.

## Data availability

NMR backbone chemical shift assignments (^1^H, ^15^N, ^13^C’, and ^13^Cα and ^13^Cβ) for synaptobrevin-2 have been deposited to the BMRB with accession code 53437. All other data presented in this study are available from the corresponding author upon request.

## Acknowledgments

We thank Dr. Buket Rüffer and Dr. Dominik Dluhosch for helpful discussions about MST measurements. N.L. acknowledges the German Science Foundation for funding through the Heisenberg Program (DFG grant number 433700474) as well as access to the Jülich-Düsseldorf Biomolecular NMR Center, jointly run by Forschungszentrum Jülich and Heinrich-Heine-University Düsseldorf (HHU).

## Authorship contribution

**Sophia Werner:** Investigation, Methodology, Formal analysis, Resources, Writing. **Tobias Stief:** Investigation, Methodology, Formal analysis, Resources, Writing. **Celina Schulz:** Investigation, Methodology, Formal analysis, Writing. **Katharina Vormann:** Resources, Writing. **Robin Backer:** Resources, Writing. **Marie P. Schützmann:** Resources. Writing. **Lothar Gremer:** Investigation, Formal Analysis, Writing. **Wolfgang Hoyer:** Investigation, Formal Analysis, Writing. **Nils-Alexander Lakomek:** Conceptualization, Methodology, Writing, Supervision, Funding acquisition.

