## Supplementary Information for "Direct NMR evidence for transient binding of the vesicular SNARE protein synaptobrevin-2/VAMP2 to the α-synuclein C-terminus"

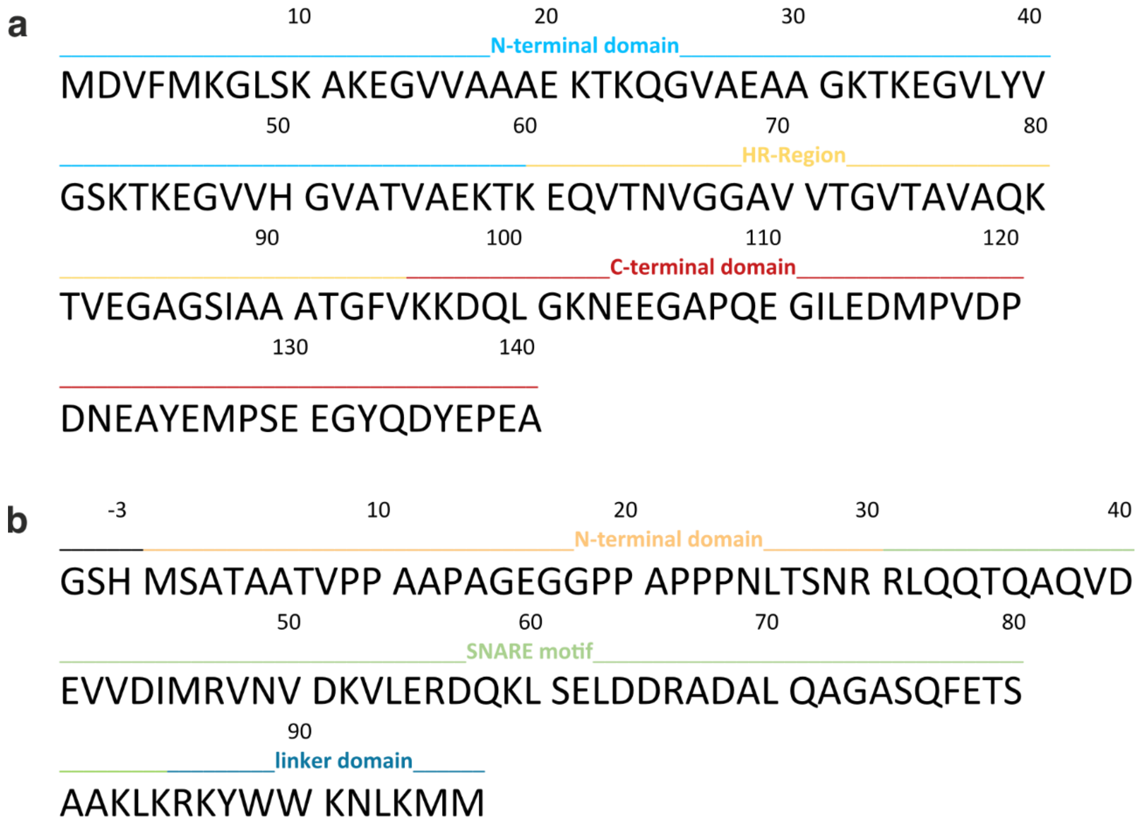


**Figure S1: Sequential information of (a) α-synuclein and (b) synaptobrevin-2(1–96).** Shown is the amino acid sequence in one-letter code. Structural components of both proteins are highlighted in different colors (see also **Figure 1**). Above the residue number is displayed. For syb-2(1–96), the three amino acids remaining from the thrombin cleavage site (GHS) are indicated by a negative sign (-3).

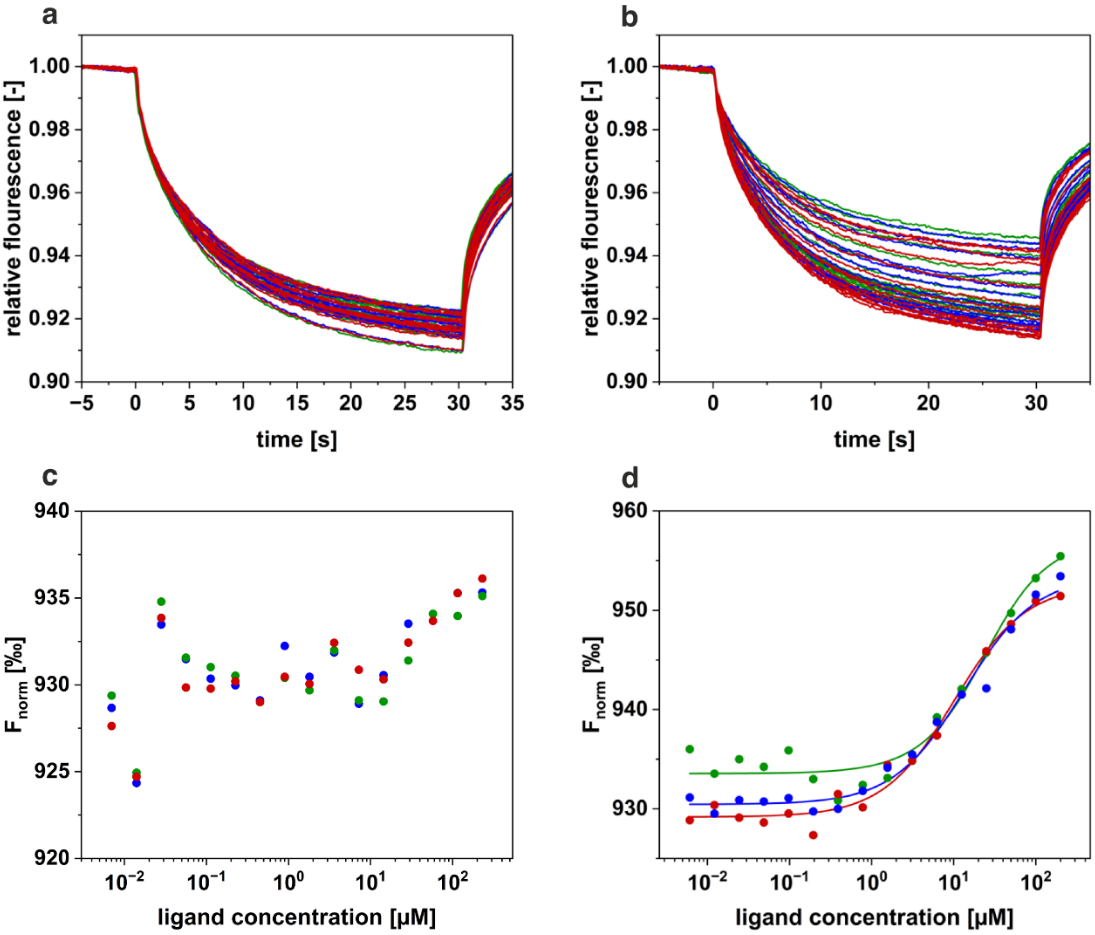


**Figure S2: Comparison of MST measurements of α-synuclein and α-synuclein$^{\Delta Q109\text{-}A140}$.** The MST traces are plotted with the relative fluorescence against the time in seconds (100 nM α-syn$^{\Delta Q109\text{-}A140}$, different concentrations of syb-2(1-96), 20 mM sodium phosphate, pH 6.0, 50 mM NaCl). All measurements were performed in triplicate as indicated by the different colors. In (**a**)**,** the MST traces of the measurement with the truncated α-syn variant are shown. (**b**) displays the MST traces of the measurement with full-length α-syn. In (**c**) (α-syn$^{\Delta Q109\text{-}A140}$) and (**d**) (α-syn full length), all $K_d$ fit data are displayed. The fitted and normalized data ($F_{norm}$ [‰]) were plotted against the concentration of the ligand syb-2(1–96) on a logarithmic scale.

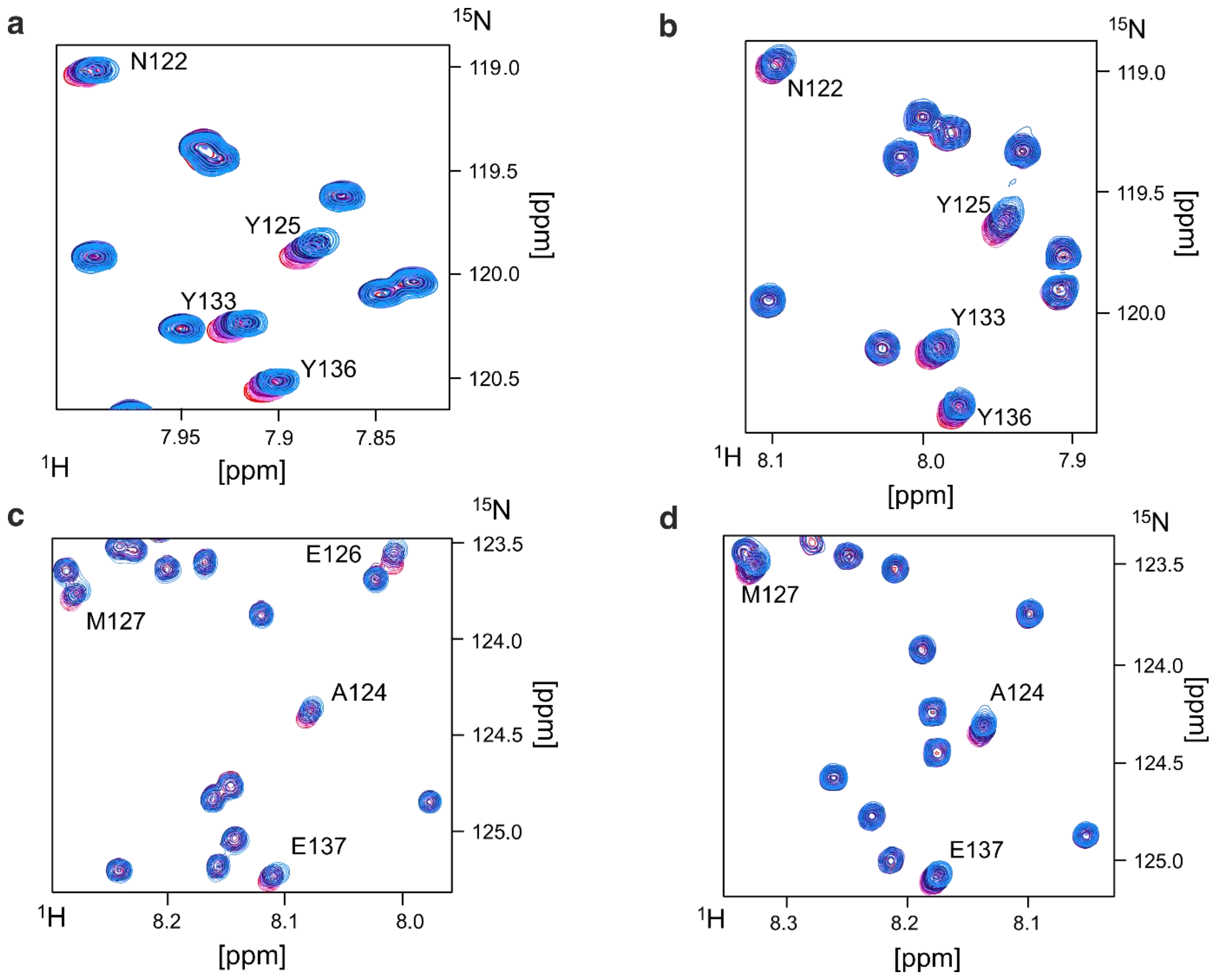


**Figure S3: NMR titration of α-synuclein with synaptobrevin-2(1–96) zoomed results.** Overlay of zoomed-in HSQC spectra of 40 µM $^{15}$N α-syn in the presence of syb-2(1–96), measured at 1.2 GHz at (**a** and **c**) 20 °C and at (**b** and **d**) 37 °C (20 mM sodium phosphate, pH 6.0, 50 mM NaCl). All titration steps are shown (reference red; molar ratio of: 1:1, pink; 1:2, light violet; 1:3, dark violet; 1:4, dark blue; 1:5, light blue).

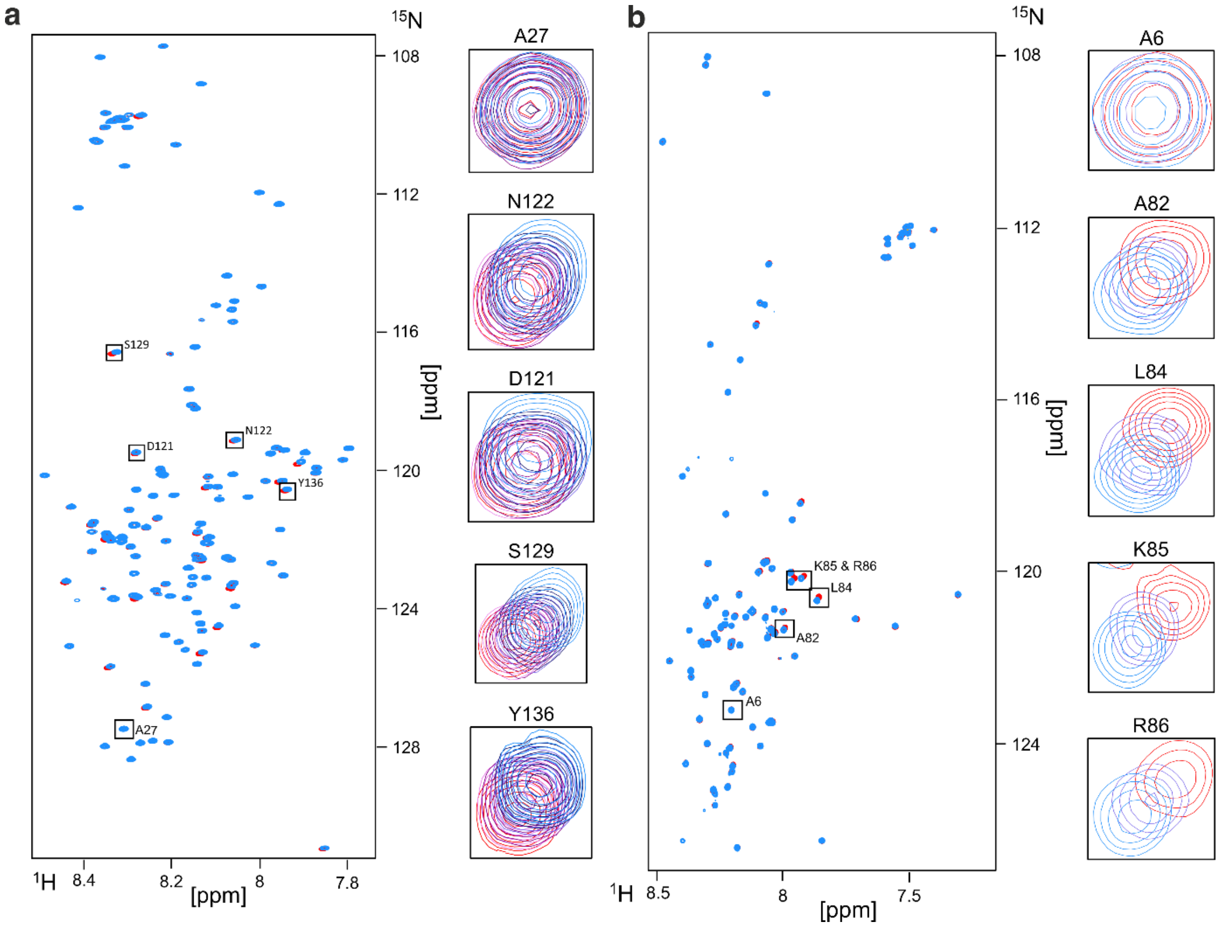


**Figure S4: NMR titration of α-synuclein with synaptobrevin-2 (1-96) and vice versa at 37 °C.** Overlay of the HSQC spectra of (**a**) 40 µM $^{15}$N α-syn in the presence of syb-2(1-96) (1:5, blue) and absence of syb-2 (1-96) (red), measured at 1.2 GHz at 37 °C. (**b**) 40 µM $^{15}$N syb-2(1-96) in the presence of α-syn (1:5, blue) and in the absence of α-syn (red), measured at 1.2 GHz at 37 °C. Some residues with large chemical shifts are shown separately with an overlay of all titration steps (reference red, molar ratio of: 1:1, pink; 1:2, light violet; 1:3, dark violet; 1:4, dark blue; 1:5, light blue). A27 and A6 are shown for reference purposes, respectively.

Table S1: Assignment of α-synuclein backbone amide [$^1$H, $^{15}$N] resonances at 20 °C and 37 °C.

Table S2: Assignment of synaptobrevin-2 backbone amide [$^1$H, $^{15}$N] resonances at 20 °C and 37 °C.
